# The Effects of Speech Masking on Neural Tracking of Acoustic and Semantic Features of Natural Speech

**DOI:** 10.1101/2023.02.10.527537

**Authors:** Sonia Yasmin, Vanessa C. Irsik, Ingrid S. Johnsrude, Björn Herrmann

## Abstract

Listening environments contain background sounds that mask speech and lead to communication challenges. Sensitivity to slow acoustic fluctuations in speech can help segregate speech from background noise. Semantic context can also facilitate speech perception in noise, for example, by enabling prediction of upcoming words. However, not much is known about how different degrees of background masking affect the neural processing of acoustic and semantic features during naturalistic speech listening. In the current electroencephalography (EEG) study, participants listened to engaging, spoken stories masked at different levels of multi-talker babble to investigate how neural activity in response to acoustic and semantic features changes with acoustic challenges, and how such effects relate to speech intelligibility. The pattern of neural response amplitudes associated with both acoustic and semantic speech features across masking levels was U-shaped, such that amplitudes were largest for moderate masking levels. This U-shape may be due to increased attentional focus when speech comprehension is challenging, but manageable. The latency of the neural responses increased linearly with increasing background masking, and neural latency change associated with acoustic processing most closely mirrored the changes in speech intelligibility. Finally, tracking responses related to semantic dissimilarity remained robust until severe speech masking (−3 dB SNR). The current study reveals that neural responses to acoustic features are highly sensitive to background masking and decreasing speech intelligibility, whereas neural responses to semantic features are relatively robust, suggesting that individuals track the meaning of the story well even in moderate background sound.

## Introduction

Many sound environments in everyday life contain background sounds, such as ambient music or speech, that can mask the target speech signal, resulting in communication challenges (Mattys, 2012; Song et al, 2011; Meyer et al., 2013). Segregation of speech from background sound is facilitated by a host of acoustic features such as onset times and harmonicity (Carroll et al., 2011; Flaherty et al., 2021; Kong et al., 2012; Darwin & Carlyon, 1995; Darwin, 2008). For example, speech signals fluctuate in amplitude at the semi-regular rate at which syllables, and words are uttered, typically below 10 Hz (Rosen, 1992). Because the amplitude fluctuations in speech and background sound typically differ, sensitive tracking of the amplitude fluctuations of speech provides a means to segregate speech from background sound. Semantic information also facilitates speech-in-noise perception. The semantic context of what has been heard can be used to predict upcoming words and, in turn, improve speech intelligibility in challenging listening conditions (Holt & Bent, 2017; Holiday et al., 2008; Shi, 2014; Zekveld et al., 2011; Davis & Johnsrude, 2007; Miller et al., 1951, Ganong, 1980; Pitt & Samuel, 1993; Norris et al., 2003). This is especially important for individuals with hearing impairments, who experience disproportionate challenges in settings with noisy backgrounds (Henry & Heinz, 2012; Monaghan et al., 2020; Bacon et al., 1998; Alain et al., 2014). Understanding how neural encoding of acoustic and semantic information occurs in different individuals and contexts is an important step towards clinical interventions for hearing loss, which are critically needed. The current study is concerned with how neural encoding of the acoustic amplitude fluctuations and the semantic context (measured based on semantic dissimilarity between words) of speech is affected by different degrees of background masking noise. This is accomplished by measuring neural tracking-responses between naturalistic stimulus properties and associated electrophysiological activity.

Much of the research into the neural processing of acoustic and semantic features of speech has relied on brief, disconnected sentences presented in a repetitive event-related design (Uhrig et al., 2020; Kasparian et al., 2016; Handy, 2005; Luck, 2014; Salmelin, 2007; Picton, 2013; Pratarelli et al., 1995; Lovrich et al., 1988; Connolly et al., 1994). However, speech in everyday life is typically more continuous (Schiffrin et al., 1984; Jefferson et al., 1978; Ochs et al., 1992; Pasupathi et al., 2002; Ochs et al., 1992), requires the integration of words into a larger semantic context and topical thread (Ehrlich and Rayner, 1981; Hale, 2001; Frank, 2013; Smith & Levy, 2013), and may be intrinsically motivating for a listener to comprehend. Listeners may thus engage differently with continuous speech compared to disconnected sentences, and the recruited neural mechanisms may thus also differ.

We have recently shown that listeners are absorbed by and enjoy spoken stories, even when they experience effort and miss occasional words as a result of moderate background masking (Herrmann & Johnsrude 2020). Engagement measured neurally through across-participant synchronization of neural activity also appears to be little affected by moderate background masking (Irsik et al., 2022a). Moreover, older adults appear to benefit from speech glimpses in background noise for comprehension more when listening to spoken stories than when listening to disconnected sentences (Irsik et al., 2022b). This suggests that something about the stories – perhaps the degree to which they pique interest and motivate listening? – is resulting in a qualitatively different listening behaviour in older people compared to disconnected sentences.

The neural processing of continuous speech is often measured by calculated a linear mapping between features of a continuous speech stimulus and the electro- or magnetoencephalographic (EEG/MEG) signals recorded while participants listen to the speech (Crosse et al., 2016; Das et al., 2020; Iotzov, & Parra, 2019; Synigal, et al., 2020). The result of such stimulus-to-neural-response mapping is the temporal response function (TRF; Crosse et al., 2016; Broderick et al., 2018; Crosse et al., 2021). TRF deflections can be interpreted similarly to components of the event-related potential for discrete speech tokens such as words (Broderick et al., 2018; Crosse & Lalor, 2014; Luck, 2012; Luck, 2014). The TRF approach has most frequently been used to investigate how acoustic properties of speech, such as the amplitude envelope, are encoded in the brain, and how this differs as a function of task demands (Das et al., 2020; Das et al. 2018; Verschueren et al., 2021; Fuglsang et al., 2017; Akram et al., 2016; Teoh et al., 2019; Drennan & Lalor, 2019). For example, the magnitude of the TRF calculated for the amplitude envelope of speech is larger for speech that is attended compared to speech that is ignored in two-talker listening contexts (Verschueren et al., 2021; Fuglsang et al., 2017; Fiedler et al., 2019; Puvvada & Simon., 2017; Brodbeck et al., 2020). The degree to which neural activity tracks the acoustic speech envelope has also been linked to speech comprehension (Verschueren et al., 2021; Decruy et al., 2019; Decruy et al., 2020).

Previous studies have revealed that the N100 response (bearing resemblance to the acoustic TRF) to noise-vocoded speech is correlated with comprehension scores (Obleser & Kotz., 2011). Similarly, acoustic envelope tracking also shows a positive relationship with intelligibility (Decruy et al., 2019; Decruy et al., 2020). However, the relationship between neural tracking of acoustic speech features and speech intelligibility may not be linear. When speech is parametrically degraded using noise-vocoding, envelope tracking indexed by the TRF exhibits a U shape: amplitude is greatest for moderate levels of degradation, and smaller both for intact and for highly degraded (1-channel vocoded) speech (Hauswald et al., 2022).

Noise-vocoding differs substantially from speech masked by babble noise. The latter resembles more closely situations that most individuals experience in everyday life, and that are reported by older individuals to be challenging and effortful (Frisina et al., 1997; Gordon-Salant, 2006). Here, we investigate whether, when speech is masked by a 12-talker background babble noise at different signal to noise ratios, envelope tracking measured as the TRF exhibits a similar inverted U-shape to that observed by Hausfeld et al (2022).

TRFs have also been used to investigate whether semantic features during continuous speech listening are encoded in the brain (Broderick et al., 2018; Gillis et al., 2021; Devaraju et al., 2021). In such studies, each word in a spoken story is represented by a high-dimensional numerical vector that captures semantic information. Words for which the corresponding vectors correlate highly are more semantically similar than words for which the vectors correlate less (Pennington et al., 2014; Mikolov et al., 2013). By calculating correlations for consecutive words within a sentence or a story, a dissimilarity score can be calculated for each word, reflecting the degree to which a word is incongruent with the preceding semantic context (Broderick et al., 2018; Broderick et al., 2020; Broderick et al., 2021). These dissimilarity scores are then used to calculate a “semantic” TRF, reflecting this incongruency, or “surprisal”; which is thought to reflect the representation of contextual information in the brain (Gillis et al, 2021; Broderick et al., 2018; Broderick et al., 2020; Broderick et al., 2021).

Similar to the acoustic TRF (Hauswald et al., 2022; Das et al., 2020; Das et al. 2018; Verschueren et al., 2021; Fuglsang et al., 2017; Akram et al., 2016; Teoh et al., 2019; Drennan & Lalor, 2019), the magnitude of the TRF calculated for the semantic dissimilarity is larger for attended compared to ignored speech (Broderick et al., 2018). However, the degree to which neural encoding of semantic dissimilarity is affected by speech masking is not clear. In previous studies, speech was masked by a single talker at one signal-to-noise ratio (SNR), and the magnitude of the semantic TRF was reduced for the unattended speaker (Broderick et al., 2018; Brodbeck et al., 2018). However, single-talker masking differs substantially from multi-talker masking (Jones & Macken, 1995; Zaglauer et al., 2017; Macken et al., 2003). A single-talker masker may not overlap spectrally very much with the target (depending on the pitch difference between the target and masker voices), it will have a highly variable envelope that will differ from that of the target, and so physical interference between target and masker will be minimal. Nevertheless, a single talker masker is potentially confusable with the target, and might be distracting (Summers & Roberts., 2020) enhancing masking efficacy. Twelve-talker babble, in contrast, is more spectrally dense, and has a flatter envelope, and thus physically interferes with (i.e., energetically masks) a single-talker target more than a single-talker masker. Furthermore, 12-talker babble does not contain intelligible word-level information. Thus, results from research using single-talker masking probably will not generalize to a situation in which multiple competing talkers are present. Indeed, recent studies have found that intelligible single-talker maskers reduce acoustic tracking of the target speech when compared to babble maskers (Song et al., 2019; Song et al., 2020).

How semantic context, or dissimilarity, encoding is affected by multi-talker background noise at different SNRs is unknown. In the current study, we use 12-talker babble noise at different SNRs to investigate how SNR affects the encoding of acoustic and semantic features of speech. Given that individuals appear highly engaged in story listening even in the presence of moderate background noise (Herrmann & Johnsrude, 2020; Irsik et al., 2022a), we expect that semantic processing, indexed by semantic dissimilarity tracking, also remains high at moderate background noise, and will only decrease for highly masked speech. Moreover, the relationship between semantic tracking and intelligibility has been scarcely explored. A few studies have investigated how the N400 response, potentially similar to the semantic TRF, is related to intelligibility. These studies have revealed a positive relationship between the N400 response and intelligibility (Broderick et al., 2018; Strauß et al., 2013; Jamison et al., 2016), suggesting that we will observe a positive relationship between semantic dissimilarity tracking and behaviourally measured intelligibility.

Neural tracking of continuous speech is often investigated using audiobook narrations (Broderick et al., 2018; Broderick et al., 2020; Broderick et al., 2021). Such materials are typically well articulated, sentences build systematically on each other, and there is a clear and well-understood grammatical framework in place (Thanh, 2015; Carter & Mncarthy, 1995). Speech in everyday life is subject to more disfluencies than audiobook narrations as speakers often use slang, filler-words, sentence fragments, corrections, unintentional pauses, and more flexible grammar (Bortfeld et al., 2001; Tree et al., 1995). It is possible that these discrepancies between naturalistic speech and audiobook narrations may affect intelligibility, effort, and/or neural processing (Arnold et al., 2003; Brennan et al., 2001). Because we are interested primarily in naturalistic listening, we use engaging, spoken stories from the story-telling podcast The Moth (https://themoth.org; Regev et al., 2019; Simony et al., 2016; Irsik et al., 2022a) which may mirror speech in everyday life more closely than do audiobooks (Ochs & Capps, 1996; Ervin-Tripp & Küntay, 1997).

In the current study, we use spoken stories to investigate how neural tracking of the acoustic amplitude fluctuations (envelopes) and semantic context of engaging, naturalistic speech are affected by background babble noise, and relate this to speech intelligibility of the same materials. We construct TRFs by linearly mapping acoustic and semantic features of speech onto corresponding EEG activity (Crosse et al., 2016; Crosse et al., 2021).

## Methods

We re-analyzed EEG and behavioural data from a previous study (Irsik et al., 2022a). With a few minor exceptions indicated explicitly below, the analyses, results, and conclusions are novel and do not overlap with those reported previously (Irsik et al., 2022a). We provide the relevant information about stimuli, procedures, and methods, and also refer to the details provided previously (Irsik et al., 2022a).

### Participants

Thirty-nine EEG datasets (mean age of participants: 20.3 years; age-range: 18-32 years; 19 males 20 females) and 82 behavioural data sets (mean age of participants: 28.8 years; age-range: 18-36 years; 51 males 31 females) were available for analysis. All participants provided informed written consent and were without self-reported hearing loss, neurological issues, or psychiatric disorders. The study was conducted in accordance with the Declaration of Helsinki, the Canadian Tri-Council Policy Statement on Ethical Conduct for Research Involving Humans (TCPS2-2014), and approved by the local Health Sciences Research Ethics Board of the University of Western Ontario (REB #112015; REB#112574).

### Acoustic stimulation and procedure

Each of the 39 participants listened to four spoken stories from The Moth podcast (https://themoth.org): *Reach for the Stars One Small Step at a Time* (by Richard Garriott, ∼13 min), *The Bounds of Comedy* (by Colm O’Regan, ∼10 min), *Nacho Challenge* (by Omar Qureshi, ∼11 min), and *Discussing Family Trees in School Can Be Dangerous* (by Paul Nurse, ∼10 min). Twelve-talker babble noise, taken from the revised Speech in Noise (R-SPIN) test (Bilger, 1984), was added to the stories at five different signal-to-noise ratios (SNRs): clear, +12, +7, +2, −3 dB. The SNR changed every 30-33 seconds to one of the five levels without repeating the same level twice in direct succession. When mixing stories with maskers, both the level of the story and the babble masker were adjusted in order to ensure sound level remained constant throughout each story, and was consistent across the stories. Stories were played via headphones (Sennheiser HD 25 Light) in a single-walled sound-attenuating booth (Eckel Industries) and participants were instructed to listen carefully to each story. After each story, participants answered ten comprehension questions about the story to ensure they were paying attention.

Speech intelligibility for each story, measured as words reported from target phrases, across different signal to noise ratios, was assessed in a separate group of 82 participants using online platforms for experiment hosting (Pavlovia) and recruitment (MTurk, CloudResearch interface). Each participant listened to the same materials described above; specifically one of four stories for which SNR changed about every 30-33 seconds to one of five levels (clear, +12, +7, +2, −3 dB). For each story, 80 or 100 phrases/sentences (4-8 words) were selected for intelligibility testing (4 phrase/sentences per 30-33s segment). During the experiment, one of the four spoken stories was played to a participant. The story paused occasionally (about every 16 s), and the participant was asked to type the last phrase/sentence uttered before the story paused into a text box. Just before the target utterance was heard, a fixation cross on the screen changed colour to tell participants that they had to remember verbatim what they were about to hear, and then changed colour again for the duration of the phrase/sentence. That target phrase/sentence was then reported during the pause that immediately followed (for details see Irsik et al., 2022a). The story then resumed from the beginning of the target utterance. Intelligibility was calculated as the proportion of correctly reported words, separately for each SNR condition.

### EEG recording and preprocessing

EEG was recorded from 64 active electrodes (Ag/AgCl) placed on the scalp using an electrode cap according to the 10/20 system (Biosemi ActiveTwo system) and both mastoids. A feedback loop between the common mode sense (CMS) active electrode and a driven passive electrode (see www.biosemi.com/faq/cms&drl.htm) was used as a reference for all other electrodes. EEG was recorded at a sampling frequency of 1024 Hz (208-Hz low-pass filter).

The data were pre-processed offline using custom MATLAB scripts and the Fieldtrip toolbox (Oostenveld et al., 2011). Data were re-referenced by subtracting the average across both mastoids from each channel. Line noise was suppressed using a 60-Hz notch filter. The data were high-pass filtered (0.5 Hz, 3429 points, Hann window) and low-pass filtered (22 Hz, 211 points, Kaiser window). Continuous EEG data were segmented into separate time series time-locked to story onset and downsampled to 256 Hz. Independent components analysis was used to remove signal components reflecting blinks, eye movement, and muscle activity (Makeig et al., 1996). Additional artifacts were removed after the independent components analysis by setting the voltage for segments in which the EEG amplitude varied more than 80 µV within a 0.2-s period in any channel to 0 µV (cf. Dmochowski et al., 2012, 2014; Cohen and Parra, 2016). As a last step prior to TRF analyses, data were low-pass filtered at 10 Hz (141 points, Kaiser window), because neural signals in the low-frequency range are most sensitive to acoustic and semantic features (Zuk et al., 2021; Di Liberto et al., 2015).

### Speech transcription and identification of word-onset times

Transcription for stories were done manually for each story. Non-words and incomprehensible mumbles were ignored for the analysis of EEG. The onset time for each word in each story was obtained using the Clarin’s forced alignment software (Schiel, 1999). Onset times were manually verified, and incorrect estimations were manually corrected.

### Acoustic and semantic temporal response functions

We used a forward model based on the linear temporal response function (TRF; Crosse et al., 2016; Crosse et al., 2021) to separately model the relationship between features of the auditory stimulus and EEG activity (see Figure 1). The TRF model uses linear regression with ridge regularization (Crosse et al., 2016; Crosse et al., 2021; Hoerl & Kennard 1970a; Hoerl & Kennard 1970b). Based on previous work, the ridge regularization parameter λ was set to 10 (Fielder et al., 2019; Fielder et l., 2017).

**Figure 1.**
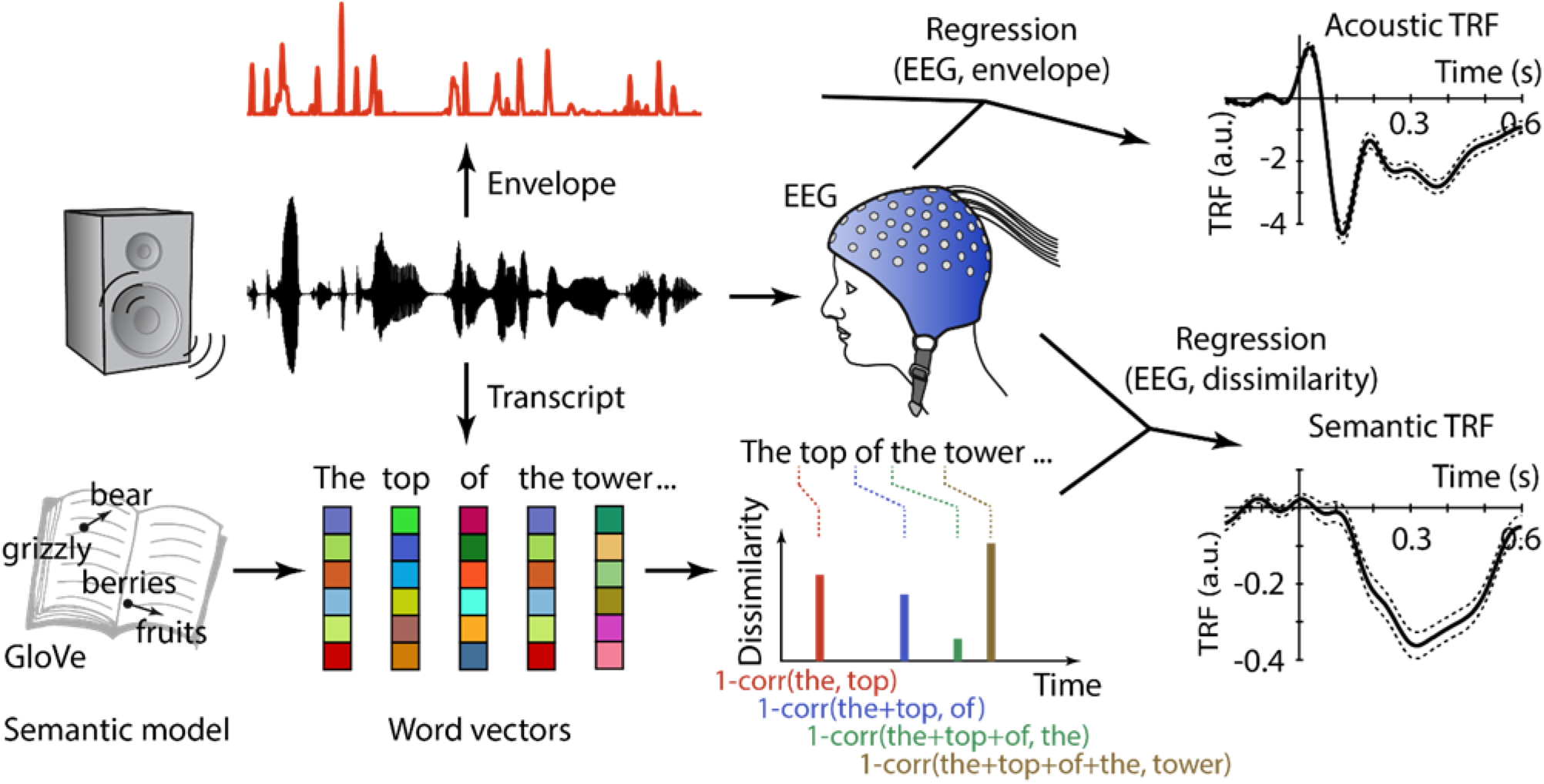
Schematic of the procedure for obtaining acoustic and semantic temporal response functions (TRFs). The middle row schematically shows stimulation, acoustic waveform, and EEG recording. Top row represents the calculation of the onset-amplitude envelope from the acoustic waveform. The amplitude envelope is regressed against the EEG data to obtain an acoustic TRF. The bottom row schematically displays the calculation of the semantic TRF. Global Vectors for Word Representation (GloVe) pretrained vectors, representing word meaning, were identified for each word of each story. Colors in the vectors schematically represent different magnitudes. A semantic dissimilarity value was calculated for each word as 1 minus the correlation between the current word’s vector and the averaged vectors across all preceding words of a sentence. A dissimilarity regressor was created by placing each word’s dissimilarity value at its respective word-onset time (while values at all other time points were zero). The dissimilarity vector is regressed against the EEG data to obtain a semantic TRF.

The current TRF analyses focused on two representations of the auditory stimulus: the cochlear envelope (i.e., envelope of a modelled cochleogram) and semantic dissimilarity. To estimate the acoustic representation for each story, we modelled the cochleogram for the acoustic waveform of each story using Lyon’s Passive Ear model (Slaney, 1988a) as implemented in the Auditory Toolbox Version 2 (Slaney, 1998b). The toolbox calculates the auditory nerve responses using the probability of firing along the auditory nerve given the acoustic properties of an input sound (Figure 1). We then averaged across all auditory filters of the cochleogram. The analytic Hilbert transform of the amplitude envelope was calculated. We low-pass filtered the envelope using a 40-Hz filter (Butterworth filter), calculated the first derivative, and set all negative values to zero in order to obtain the amplitude-onset envelope (Fiedler et al., 2017; Fiedler et al., 2010; Hertrich et al., 2012). This amplitude-onset envelope was used as a regressor for the TRF analysis (Figure 1).

Representations for semantic dissimilarity were obtained using previously described methods (Broderick et al., 2018; Broderick et al., 2018). We utilized pretrained vectors from the Global Vectors for Word Representation (GloVe) project to obtain a semantic representation for each word in form of numerical vectors (i.e., word embeddings; 300 dimensions; Pennington et al., 2014; https://nlp.stanford.edu/projects/glove/). GloVe is an unsupervised learning model that maps words into vector space based on their semantic relationships. The numerical vectors of words that are semantically more similar are more correlated (e.g., frog vs toad) compared to the vectors of words that are semantically less similar (e.g., frog vs shoe). The GloVe corpus consists of 400,000 vocabulary entries and their corresponding numerical vectors. For each word of the story transcripts, we obtained the corresponding word vector from GloVe, if it existed in the corpus. On average across the four stories, 11% of words were not available in the GloVe corpus and they were thus not considered for calculating semantic dissimilarity regressors for the EEG TRF analysis. Using the word vectors, a semantic dissimilarity value was calculated for each word of each story based on the local sentence context (Broderick et al., 2018). Specifically, the Pearson correlation between the vector of the current word and the averaged vectors across all preceding words of the sentence was calculated. Each correlation value was subtracted from 1 to calculate the dissimilarity value (Figure 1). A regressor for the TRF analysis was then created by placing each word’s dissimilarity value at its respective word-onset time (while values at all other time points were zero). This regressor was created at the sampling frequency of the EEG data.

Because the dissimilarity regressor contains impulses at word onsets (with values being otherwise zero), it is sensitive to brain responses associated with the acoustic onset of words. In order to mitigate the influence of acoustic properties on the semantic TRF, we also calculated a ‘static’ TRF, where the regressor is calculated using the median dissimilarity value across words for all word onsets (all other samples remain zero). Hence, this regressor also contained impulses at word onsets, but the impulses were all of similar magnitude and no semantic dissimilarity variations were represented.

For each participant, EEG channel, and ∼30 s data segment corresponding to different SNR levels within stories, a TRF was calculated using time windows of −0.3 s to 0.7 s between the input time series of stimulus features (acoustic, semantic) and the corresponding EEG time courses, measured from word onset. TRFs were averaged across segments (approximately 30 s), separately for each SNR level. To obtain the final semantic TRF, we subtracted the ‘static’ TRF from the TRF derived using the dissimilarity vector. The result of these TRF calculations was one acoustic TRF and one semantic TRF for each SNR level, EEG channel, and participant.

### Analysis of the relation between SNR levels and TRF amplitude and latency

For the analysis of amplitude and latency of specific deflections in the TRF, we averaged across a subset of fronto-centro-parietal channels (FC1, FC2, FCz, FC3, FC4, C1, C2, Cz, C3, C4, CP1, CP2, CPz, CP3, CP4, P1, P2, Pz, P3, P4) known to be sensitive to responses elicited by acoustic and semantic manipulations (Broderick et al., 2018; Connolly et al., 1992; Connolly et al., 1994; Martin et al., 1999; Finke et al., 2016; Martin et al., 2005). We used custom MATLAB scripts to automatically identify response peaks within selected time ranges. For the acoustic TRF, we estimated the peak latency for the negative deflection within 100-250 ms for each participant and SNR level. We call this negative deflection the “acoustic tracking response”. Although there is obvious resemblance to the typical N1/N100 component of event-related potentials (Crosse & Lalor, 2014), we want to avoid the assumption that what we observe here is indeed the N1/N100. The amplitude for the acoustic tracking response was calculated as the mean amplitude across 10 ms centered on a participant’s individual peak latency. Our investigations for the acoustic TRF are restricted to the negativity at 100-250 ms, as visual inspection of the time course in Figure 2a demonstrates this peak to be most susceptible to SNR-related changes.

**Figure 2.**
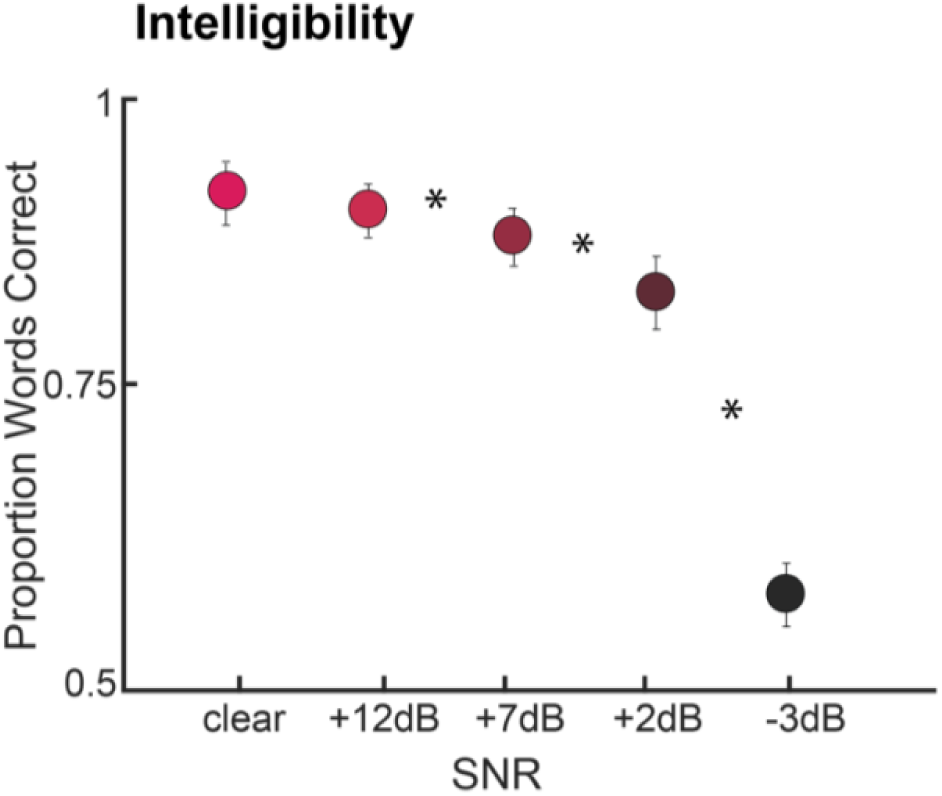
Effects of SNR on Intelligibility. Mean proportion of correctly reported words plotted as a function of SNR (clear, +12, +7, +2, −3 dB SNR). Asterisks indicate that the intelligibility of the two flanking SNRs differ significantly. Error bars reflect the standard error of the mean. *p < 0.05. For more detailed information see Irsik et al., 2022a.

For the semantic TRF, we estimated the peak latency for the negative deflection within 300-450 ms for each participant and SNR level. We call this negative deflection the “semantic tracking response”. This deflection resembles the typical N400 component of event-related potentials, which has been associated with semantic incongruency (Kutas & Federmeier, 2011; Broderick et al., 2018), but, again, we do not assume that what we observe here is indeed the N400. The amplitude for the semantic tracking response was calculated as the mean across 100 ms centered on a participant’s individual peak latency.

We evaluate the degree to which acoustic and semantic tracking changes linearly or quadratically over SNRs. To this end, a quadratic function was fitted separately to the latency and amplitude data as a function of SNR levels (coded: [-2 −1 0 1 2]), separately for each participant. Quadratic fits, appropriate to test whether the data conform to a U-shape, as predicted, were calculated separately for the acoustic TRF (acoustic tracking response) and the semantic TRF (semantic tracking response), and separately for both amplitude and latency data. The resulting linear and quadratic coefficients were tested against zero using a one-sample t-test to identify significant nonzero linear and quadratic trends of TRF amplitude/latency as a function of SNR.

We also conducted one-way repeated measures ANOVAs (rmANOVAs) to quantify effects of SNR on acoustic and semantic TRF amplitudes and latencies. We performed posthoc pairwise comparisons using independent samples t-tests, with false discovery rate correction (FDR; Benjamini and Hochberg, 2016), between neighboring SNR levels to evaluate differences. FDR corrected p-values are referred to as *p_FDR_*.

### Relationship between acoustic and semantic TRFs, and speech intelligibility

Amplitudes and latencies of acoustic and semantic TRFs as well as speech intelligibility (from online testing; Figure 2) have different units and magnitudes. In order to compare them directly, we calculated z-scores for each participant. That is, separately for each individual and dependent measure, we took the value at each SNR, subtracted the average across the five SNRs, and then divided by the standard deviation of that measure across SNRs. Z-score normalized acoustic TRF amplitude and latency, and z-normalized semantic TRF amplitude and latency were also sign inverted by multiplying the data by −1, to ensure that larger values indicate larger amplitudes and shorter latencies, enabling comparison with speech intelligibility data (for which a larger value means better comprehension). In order to compare these responses, we again fit quadratic functions separately to the acoustic TRF amplitude, semantic TRF amplitude, acoustic TRF latency, and semantic TRF latency, and to the speech intelligibility data, across SNRs. We used t-tests on the resulting coefficients to examine whether changes across SNR in speech intelligibility were more similar to the acoustic TRF, the semantic TRF, or neither.

### Effect size

Effect sizes are reported as partial eta squared for ANOVAs (η^2^_p;_ Kennedy, 1970) and Cohen’s d for *t*- tests (d; Cohen., 1988).

## Results

### Amplitude and latency of acoustic TRFs are modulated by the degree of background masking

We found that the amplitude of the acoustic tracking response was quadratically modulated by SNR (t_38_ = 9.225, p = 3.06 × 10^-11^, d = 1.477). There was no significant linear modulation of acoustic tracking response amplitude by SNR (t_38_ = −1.556, p = 0.1281, d = 0.249). To further explore the quadratic effect, we conducted a rmANOVA (F_4,152_ = 19.537, p = 5.44 × 10^-13^, η^2^_p_ = 0.3396), followed by pair-wise comparisons between SNR levels. After false discovery rate (FDR) correction, we observed significant differences for all neighboring SNR levels except between the +12 dB to +7dB conditions (clear smaller than +12 dB: t_38_ = 6.194, p_FDR_ = 3.1 × 10^-7^, d = 1.099; +12 vs +7 dB: t_38_ = 1.866, p_FDR_ = 0.069, d = 0.333; +7 greater than +2 dB: t_38_ = 3.262, p_FDR_ = 0.0023, d = 0.607; +2 greater than –3 dB: t_38_ = 2.423, p_FDR_ = 0.0203, d = 0.460). These results indicate a U-shape, as predicted: the acoustic tracking response amplitude increased for minimal to moderate background noise relative to clear speech, and then decreased again for speech that is highly masked (Figure 3B).

**Figure 3.**
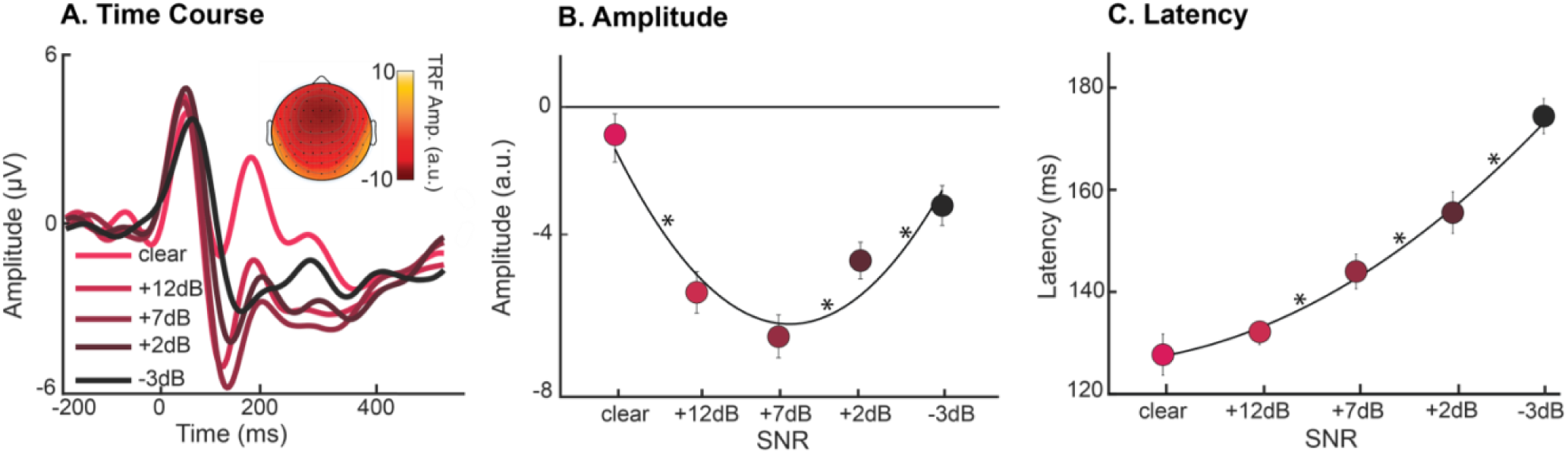
Effects of SNR on acoustic TRFs. **A.** TRF time courses (averaged across fronto-central-parietal electrode cluster) for each SNR condition and scalp topography for the acoustic tracking response (negative deflection at around 150 ms). **B.** The mean acoustic tracking response amplitude across participants, displayed for each SNR condition. Significant differences in response magnitude exist between clear and +12 dB SNR, +7 dB and +2 dB SNR, and +2 dB and −3 dB SNR. **C.** The mean acoustic tracking response latency across participants, displayed for each SNR. Significant differences in response latency exist between +12 dB and +7 dB, +7 dB and +2 dB, and +2 dB and −3 dB. The black lines in panels B and C indicate the best fitting line from a quadratic fit. Error bars reflect the standard error of the mean. *neighbouring SNRs differ at p < 0.05.

Acoustic tracking response latency increased linearly with decreasing SNR (t_38_ = 10.979, p = 2.39 × 10^-13^, d = 1.758). There was also a quadratic relationship between SNR and acoustic tracking response latency (t_38_ = 2.452, p = 0.0189, d = 0.393). We followed up on the linear and quadratic effects with an rmANOVA (F_4,152_ = 43.61, p = 2.5 × 10^-10^, η^2^_p=_ 0.534) and pair-wise comparisons between neighboring SNR levels. After FDR correction, all neighboring SNR levels differed significantly except the clear and +12 dB conditions (clear vs +12 dB: t_38_ = −1.254, p_FDR_ = 0.218, d = 0.209; +12 vs +7 dB: t_38_ = 3.880, p_FDR_ = 0.0004, d = 0.667; +7 vs +2 dB: t_38_ = 3.355, p_FDR_ = 0.0018, d = 0.493; +2 vs – 3 dB: t_38_ = 4.183, p_FDR_ = 0.0002, d = 0.802).

### Amplitude and latency of semantic TRFs are modulated by the degree of background masking

We evaluated the relationship between the degree of background masking of speech and the neural responses to semantic encoding of the story (i.e., semantic dissimilarity; Figure 4). We observed that the semantic tracking response amplitude was quadratically modulated by SNR (t_38_ = 2.731, p = 0.0095, d = 0.437), whereas the linear modulation was not significant (t_38_ = 0.872, p = 0.389, d = 0.1397). We followed up on this result using an rmANOVA (F_4,152_ = 2.706, p = 0.032, η^2^_p_ = 0.0665) and pair-wise comparisons between neighboring SNR levels. After FDR correction, the semantic tracking response amplitude was lower at the least favourable SNR condition compared to its neighbour (−3 dB and +2 dB SNR; t_38_ = 3.399, p_FDR_ = 0.0016, d = 0.542), whereas tracking did not differ between any other pairs (for all p_FDR_ > 0.05).

**Figure 4.**
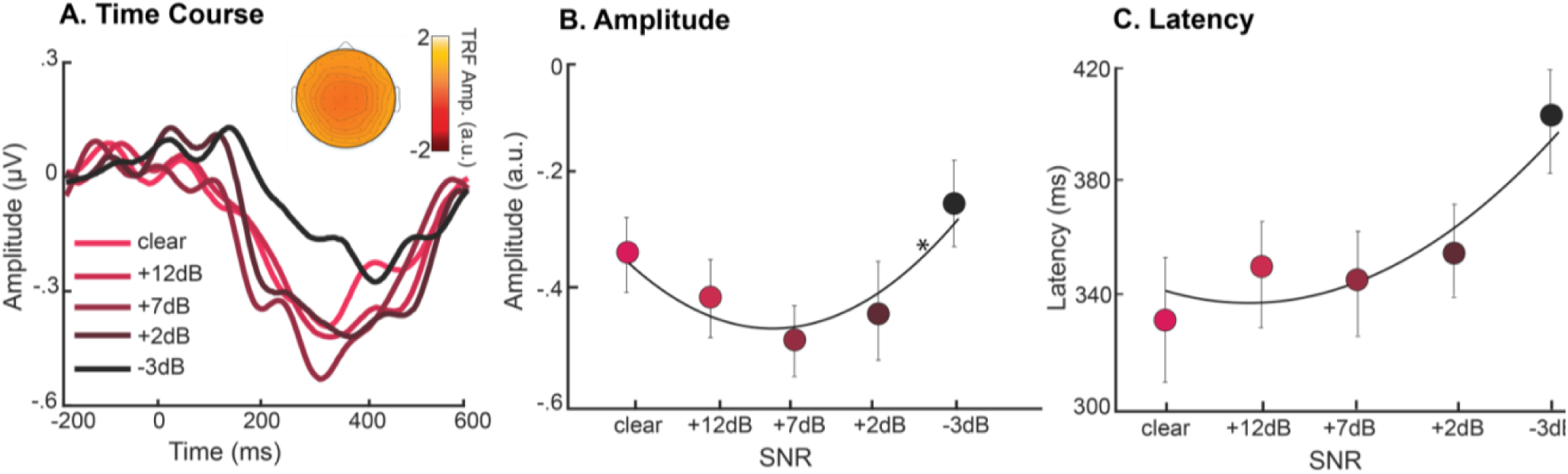
Effects of SNR on semantic TRFs. TRF time courses (averaged across fronto-central-parietal electrode cluster) for each SNR condition and scalp topography for the semantic tracking response (negative deflection at around 350 ms). **B.** The mean semantic tracking response amplitude across participants, displayed for each SNR condition. Significant differences in response amplitude exist between the +2 dB and −3 dB conditions, only. **C.** The mean semantic tracking response latency across participants, displayed for each SNR. No significant differences in response latency exist between neighboring conditions. The black lines in panels B and C indicate the best fitting line from a quadratic fit. Error bars reflect the standard error of the mean. *p < 0.05.

As for the acoustic tracking response, the semantic tracking response latency increased linearly with decreasing SNR (t_38_ = 2.834, p = 0.0073, d = 0.454), and no quadratic trend was found (t_38_ = 1.211, p = 0.233, d = 0.194). The rmANOVA revealed a significant effect of SNR (F_4,152_ = 3.043, p = 0.019, η^2^_p_ = 0.074), although no two SNR levels differed after FDR correction (neighboring and not).

### Comparison of semantic and acoustic TRFs and their relation to speech intelligibility

In order to investigate differences in how SNR affected neural acoustic and semantic tracking, and to examine whether the change in intelligibility over SNR related to the acoustically driven responses or the semantically driven responses, quadratic functions were fit to z-scored data and the resulting linear and quadratic coefficients were compared between measures. We first contrasted coefficients between the acoustic and the semantic tracking responses, before comparing each of these to coefficients from fits to intelligibility data.

The acoustic tracking response amplitude showed a stronger linear relationship with SNR (positive relationship) than the semantic tracking response amplitude (negative relationship) (t_38_ = 2.723, p = 0.0096, d = 0.610 (Figure 5). The acoustic tracking response amplitude was also more quadratically related to SNR than the semantic tracking response amplitude (t_38_ = 4.214, p = 1.5 × 10^-4^, d = 0.962; Figure 5). This is consistent with the observation that the semantic tracking response amplitude only dropped at the lowest SNR level (−3 dB SNR; Figure 3B). As can be seen in Figure 5A, SNR had a much bigger impact on the acoustic than on the semantic tracking response amplitude.

**Figure 5.**
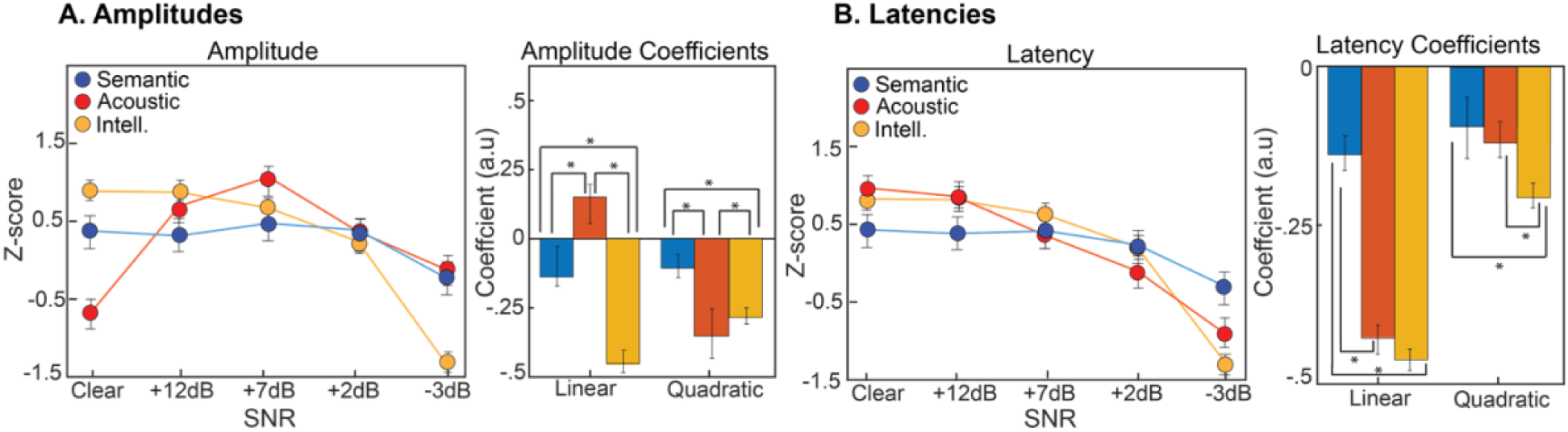
Normalized acoustic, semantic, and intelligibility data. **A left:** The mean z-scored amplitude for the acoustic tracking response and semantic tracking response (sign-inverted such that larger values mean larger responses), as well as intelligibility data are shown as a function of SNR. **Right:** The quadratic and linear coefficients obtained by fitting 2^nd^ order polynomial functions to the amplitude and intelligibility data. **B left:** The mean z-scored latency for the acoustic tracking response and semantic tracking response (sign-inverted such that larger values mean shorter latencies), as well as intelligibility data are shown as a function of SNR. **Right:** The quadratic and linear coefficients obtained by fitting 2^nd^ order polynomial functions to the amplitude and intelligibility data. Note that the behavioural intelligibility data and coefficients (in yellow) are identical between panels A and B. Error bars reflect the standard error of the mean. *p<0.05

The acoustic tracking response latency was more strongly linearly related to SNR than the semantic tracking response latency (t_38_ = 4.764, p = 2.77 × 10^-5^, d = 1.0821). Figure 5B shows that the acoustic tracking response latency strongly increases with decreasing SNR, whereas the semantic tracking response latency is less affected; this can also be seen in Figures 3 and 4. There was no difference between acoustic and semantic tracking in the degree of quadratic fit (t_38_ = −0.193, p = 0.848, d = 0.464).

In order to compare how SNR affects speech intelligibility and neural responses, we compared coefficients obtained from linear and quadratic fits (Figure 5). We found that speech intelligibility declined more linearly with decreasing SNR than did either the acoustic tracking response amplitude (t_38_ = 14.052, p_FDR_ = 4.89 × 10^-27^, d = 2.733) or the semantic tracking response amplitude (t_38_ = 8.845, p_FDR_ = 1 × 10^-14^, d =1.721). Speech intelligibility was also more quadratically modulated by SNR than the semantic tracking response amplitude (t_38_ = −3.433, p_FDR_ = 8.2 × 10^-4^, d = 0.668), but less quadratically modulated than the acoustic tracking response amplitude (t_38_ = −3.823, p_FDR_ = 2.1 × 10^-4^, d = 0.744). This is probably because the acoustic TRF magnitude increased significantly for intermediate SNRs, whereas intelligibility did not, and intelligibility appears to drop more precipitously at the lowest SNR (−3 dB) than does semantic tracking. These results indicate that the relationship between SNR and speech intelligibility is not entirely reflected either in the relationship between SNR and acoustically driven TRF amplitudes, or in the relationship between SNR and semantically driven TRF amplitudes.

There was no difference in linear coefficients between the acoustic tracking response latency (t_38_ = 0.439, p_FDR_ = 0.661, d = 0.0855) as a function of SNR, and speech intelligibility as a function of SNR. This suggests that with decreasing SNR, the linear decrease in speech intelligibility was similar in degree to the linear latency increase of the acoustic TRF. However, speech intelligibility was more quadratically modulated by SNR than was the acoustic tracking response latency (t_38_ = −4.297, p_FDR_ = 3.6 × 10^-5^, d = 0.835), likely as a consequence of a substantial drop in intelligibility for the most difficult SNR (−3 dB) that was absent for the acoustic tracking response latency. Compared to the semantic tracking response latency, speech intelligibility declined more linearly with decreasing SNR (t_38_ = 7.386, p_FDR_ = 2.3 × 10^-11^, d = 1.437) and was more quadratically modulated by SNR (t_38_ = 3.879, p_FDR_ = 1.7 × 10^-4^, d = 0.755).

The comparisons described in this section suggest that speech intelligibility is affected differently by SNR compared to acoustic and semantic TRFs. The acoustic TRF latency somewhat resembled the speech intelligibility data, although the decline in intelligibility for the least favourable SNR (−3 dB) was not matched by a corresponding latency increase in the acoustic TRF. Changes in SNR did not appear to influence the semantic TRF amplitude and latency, except at the least favourable SNR. This pattern is different to that for speech intelligibility.

## Discussion

In the current study, we investigated how the neural encoding of the acoustic envelopes and semantics of engaging, spoken stories is affected by different degrees of masking with multi-talker babble. We further examined how the effects of masker level on neural tracking relates its effects on intelligibility of the same materials. We looked for particular components, or deflections, of the acoustic and semantic tracking functions, thought to reflect acoustic and semantic processing. We observed these characteristic deflections around 100 ms for the acoustic envelope, and around 350 ms, for semantic dissimilarity, consistent with previous reports. We found that the neural tracking of the acoustic and semantic features of speech are modulated by background noise in different ways. Specifically, the amplitude of acoustic envelope tracking followed a U-shape with decreasing SNR, similar to what has been observed before (Hauswald et al., 2022). In contrast, semantic TRF amplitude was relatively stable across SNRs, dropping only at the least favourable SNR. Latencies increased linearly with decreasing SNR. The distinction between linear and quadratic relationships in these responses highlights the importance of examining a broad range of SNRs. Decreases in speech intelligibility with decreasing SNR appear to most closely resemble acoustic TRF latencies, but the profile of intelligibility across SNR otherwise did not seem to entirely reflect either acoustic or semantic processing. The current data suggest complex relationships between neural encoding of acoustic and semantic features of speech and speech intelligibility.

### Acoustic TRF is modulated by the degree of background masking

In the current study, we observed that amplitude of the neural tracking of the speech envelope was larger at moderate SNRs than for clear speech or for less favourable SNRs (Figure 3B; 5A). In contrast, the latency for the acoustic tracking response increased linearly with masking level (Figure 3C; 5B). Previous investigations using simple speech stimuli, such as “ba” and “da” sounds, masked by broadband noise, have generally observed linear reductions in response amplitude (Martin et al., 1999; Martin et al., 2005) and linear increases in response latencies with decreasing SNR (Martin et al., 1999; Finke et al., 2016; Martin et al., 2005). The latter we also observed here. Mirroring the observations for simple sounds, a few works using more complex speech stimuli have shown a larger magnitude of the acoustic TRF (Wang et al., 2020) and an increase in response latencies in the presence of competing speech, when compared to unmasked speech (Brodbeck et al., 2020).

Other recent work suggests a U-shaped relationship between neural tracking of the speech envelope and the degree of speech degradation (Hauswald et al., 2022), similar to the current study. Hauswald et al (2022) observed that the magnitude of the acoustically derived TRF was quadratically modulated such that acoustic tracking was largest for moderate levels of noise-vocoded speech, but smaller for both clear and highly degraded noise-vocoded speech (1-channel). Hauswald et al (2022) suggests that this quadratic relation may be explained by increased attention/cognitive control associated with listening effort for moderate degradation levels (cf. Pichora-Fuller et al 2016; Herrmann & Johnsrude 2020; Yerkes et al., 1908; Brehm & Self, 1989; Eckert et al., 2016; Kuchinsky et al., 2016), whereas less attention/cognitive control is deployed for highly intelligible speech and speech for which comprehension is too difficult (Hauswald et al., 2022). The fact the response amplitude elicited by simple sounds, such as tones, linearly decreases with increasing masking level (Michalewski et al., 2009; Martin et al., 1999; Martin et al., 2005) suggests that the quadratic relation observed for speech may be related to factors beyond pure acoustic processing, possibly attention/cognitive control. Indeed, neural tracking of the amplitude envelope of speech is larger for attended speech compared to ignored speech in two-talker listening contexts (Verschueren et al., 2021; Fuglsang et al., 2017). The U-shaped modulation of the acoustic TRF amplitude may thus reflect increased attention or cognitive control for moderately masked, still intelligible, speech relative to clear speech, whereas neural tracking is reduced when masking reduces speech intelligibility beyond some point, and the listener essentially ‘gives up’ (Picou & Ricketts, 2018; Pichora-Fuller et al. 2016). The response latency of the acoustic TRF, which increased linearly with increasing masker level, may reflect the acoustic impact of speech masking on envelope tracking more directly.

### Semantic TRF is modulated by masker level

We observed a negative deflection at around 300–450 ms after word onset that was associated with variations in how well a word was predicted based on semantic dissimilarity (Figure 4). This is consistent with the original work using TRFs to investigate neural processing of semantic context in continuous speech (Broderick et al., 2018; Brodereick et al., 2020; Broderick et al., 2021). This negative deflection in the TRF is also consistent with the N400 component of the event-related potential elicited by semantically incongruent words in simple sentences (Ritter et al., 1980; 2018; Nigam et al., 1992; Deacon et al., 1995; Strauß et al., 2013).

The magnitude of the semantic tracking response was similar to that for clear speech across increasing levels of speech masking, although it declined abruptly for the least favourable −3dB SNR condition, at which speech intelligibility was at around 55% (Figure 2). This pattern of stable responding with an abrupt decline is reflected in the fit of a quadratic but not linear function to the data. We also observed a trend towards increasing response latency with decreasing SNR, although this effect was weak. Previous work has demonstrated that the semantic TRF response is larger for attended compared to ignored speech when it is masked by a competing talker (Broderick et al., 2018). Noise vocoding is known to influence the magnitude and latency of the N400 response (Strauß et al., 2013), and others have demonstrated that the latency of the N400 increases when speech is masked with a babble noise (Connolly et al., 1992). Our work suggests that the semantic TRF response is relatively robust to changes in babble-noise level as long as something over 50%, but under 80%, of words are intelligible during story listening (the 5-dB resolution between SNR levels in our work does not allow a more fine-grained conclusion). It thus appears that the brain tracks semantic context well even in the presence of moderate background noise, potentially explaining why behavioural (Herrmann & Johnsrude, 2020) and neural (Irsik et al., 2022a) engagement with stories is relatively unaffected by background noise.

### Changes in speech intelligibility most closely resemble changes in acoustic response latency

Previous studies have demonstrated that the N100 (acoustic) response to noise-vocoded speech correlates with comprehension scores (Obleser & Kotz., 2011). Acoustic envelope tracking has also been shown to increase with speech understanding (Decruy et al., 2019; Decruy et al., 2020). Surprisingly, envelope tracking is larger in older compared to younger adults (Presacco et al., 2016; Presacco et al., 2019), despite the fact that older adults typically comprehend speech less well. Semantic processing, as captured by the N400 response, is also sensitive to whether or not speech was understood (Broderick et al., 2018; Strauß et al., 2013; Jamison et al., 2016). We investigated whether speech intelligibility is reflected in responses either to the acoustic or the semantic features of speech by examining function fits to intelligibility data, and to acoustic and semantic tracking amplitudes and latencies, as a function of SNR (Figure 5).

The U-shape of the acoustic TRF amplitudes over SNRs did not resemble the intelligibility data. The increase in acoustic TRF latency over SNRs was a closer match to the intelligibility data, but intelligibility appeared to decline less steeply than acoustic latency increased from clear speech to −3dB SNR (Figure 5). In contrast to the decline in intelligibility from clear speech to −3 dB SNR, the semantic TRF was robust across moderate masking levels (up to and including the penultimate masker level, +2 dB).

The current intelligibility data reflect the proportion of correctly reported words (Figure 2). Word report is an artificial task and is not identical to speech comprehension. It is possible that, if we had measured comprehension as gist report (“do you understand the utterance, yes/no”), instead of word report, we may have seen a closer correspondence to the amplitudes of the semantic tracking response. We examined the effect of a broad range of SNRs on neural tracking responses to acoustic and semantic properties of natural speech, which has previously not been explored fully. Our data suggest a complex relationship between intelligibility measured using word report, and neural tracking of different features of speech, over a range of masking levels. We see key differences in the way acoustics and semantics are tracked as a function of noise level; specifically, we observed that neural tracking of semantic dissimilarity, and thus context, is more resilient, when compared to acoustics and intelligibility, to challenging listening conditions, at least in healthy young adults.

## Conclusion

In the current study, we investigated how the EEG signal tracks the amplitude envelope and the semantic content of engaging, continuous speech, and how neural tracking is affected by different degrees of multi-talker masking. We also investigated how the effect of masking level on neural tracking related to the effect of masking level on intelligibility, measured as word report for the same story materials. The amplitude of the acoustic response was substantially greater at moderate masking levels compared either to clear speech, or to the lowest SNR, perhaps due to increased attention/increased cognitive control when speech comprehension was challenging, but manageable. In contrast, neural tracking of the semantic information was stable and robust to noise, declining only at the least favourable SNR. Response latencies increased linearly with increasing masking, more for acoustic envelope tracking than for semantic tracking. Changes in speech intelligibility with increased speech masking mirrored most closely the changes in the response latency to the acoustic envelope of speech, but were also somewhat robust to changes in SNR, averaging between 80 and 90% words reported correctly up to the least favourable SNR, where word report dropped to 50%. This stability with an abrupt decline at the lowest SNR resembles the magnitude of the neural tracking response to semantic information. Our data demonstrate how different features of the same speech signal are reflected in different aspects of the neural tracking response, measured using EEG, and point to a complex relationship between speech intelligibility and neural speech encoding.

## Acknowledgements

This research was supported by the Canadian Institutes of Health Research (MOP133450 to I.S. Johnsrude). BH was supported by the Canada Research Chair program (232733). SY was supported by the Natural Sciences and Engineering Council of Canada CGS-D scholarship and the Vector Institute Post Graduate Affiliates Program.

